# Microbial diversity and metabolic potential in cyanotoxin producing cyanobacterial mats throughout a river network

**DOI:** 10.1101/294421

**Authors:** Keith Bouma-Gregson, Matthew R. Olm, Alexander J. Probst, Karthik Anantharaman, Mary E. Power, Jillian F. Banfield

## Abstract

Microbial mats formed by Cyanobacteria of the genus *Phormidium* produce the neurotoxin anatoxin-a that has been linked to animal deaths. Blooms of planktonic Cyanobacteria have long been of concern in lakes, but recognition of potential harmful impacts of riverine benthic cyanobacterial mats is more recent. Consequently little is known about the diversity of the biosynthetic capacities of cyanobacterial species and associated microbes in mats throughout river networks. Here we performed metagenomic sequencing for 22 *Phormidium*-dominated microbial mats collected across the Eel River network in Northern California to investigate cyanobacterial and co-occurring microbial assemblage diversity, probe their metabolic potential and evaluate their capacities for toxin production. We genomically defined four Cyanobacterial species clusters that occur throughout the river network, three of which have not been described previously. From the genomes of seven strains from one species group we describe the first anatoxin-a operon from the genus *Phormidium*. Community composition within the mat appears to be associated with the presence of Cyanobacteria capable of producing anatoxin-a. Bacteroidetes, Proteobacteria, and novel Verrucomicrobia dominated the microbial assemblages. Interestingly, some mats also contained organisms from candidate phyla such as *Canditatus* Kapabacteria, as well as Absconditabacteria (SR1), Parcubacteria (OD1) and Peregrinibacteria (PER) within the Candidate Phyla Radiation. Oxygenic photosynthesis and carbon respiration were the most common metabolisms detected in mats but other metabolic capacities include aerobic anoxygenic photosynthesis, sulfur compound oxidation and breakdown of urea. The results reveal the diversity of metabolisms fueling the growth of mats, and a relationship between microbial assemblage composition and the distribution of anatoxin-a producing cyanobacteria within freshwater *Phormidium* mats in river networks.

## Introduction

When Cyanobacteria proliferate in freshwater environments, the production of toxins can threaten water quality and public health (Paerl and Otten, 2013). Harmful cyanobacterial blooms in lakes have been described for decades (Francis, 1878). In rivers, however, research on toxigenic benthic cyanobacterial mats is relatively recent (Quiblier *et al.*, 2013). Nevertheless, nuisance benthic cyanobacterial mats in rivers have already been documented across the globe, including New Zealand (McAllister *et al.*, 2016), California (Fetscher *et al.*, 2015; Bouma-Gregson *et al.*, 2017), France (Gugger *et al.*, 2005), and Spain (Cantoral Uriza *et al.*, 2017). Benthic mats are often formed by filamentous genera such as *Anabaena*, *Phormidium*, *Nodularia*, *Lyngbya*, or *Oscillatoria*, and are able to produce cyanotoxins such as anatoxin-a, microcystins, and lyngbyatoxin (Quiblier *et al.*, 2013). Given predictions of increasing cyanobacterial blooms in lakes and estuaries due to eutrophication and warming (Paerl and Huisman, 2009; Rigosi *et al.*, 2014), a better understanding of riverine benthic cyanobacterial mats is needed to anticipate environmental and ecological factors that may stimulate toxigenic benthic cyanobacterial blooms in rivers.

Heterotrophic bacteria often grow attached to, or in close proximity, to cyanobacterial filaments in benthic mats, exchanging nutrients and carbon (Paerl, 1996; Xie *et al.*, 2016). Cyanobacterial growth rates and other physiological process are often enhanced in the presence of co-occurring microbes (Allen, 1952), and few antagonisms between cyanobacteria and other bacteria have been reported (Paerl, 1996; Berg *et al.*, 2009). The microbial assemblage of lakes shifts when cyanobacteria bloom (Woodhouse *et al.*, 2016; Tromas *et al.*, 2017), and the assemblage composition differs depending on the cyanobacterial species causing the bloom (Louati *et al.*, 2015; Bagatini *et al.*, 2014). With most interactions among Cyanobacteria and co-occurring bacteria facilitating cyanobacterial growth, identifying the diversity and metabolisms of the whole microbial consortia within toxigenic cyanobacterial mats is central to understanding the ecological processes that drive the proliferation of toxigenic mats in rivers.

Little is known about the microbial consortia associated with freshwater toxigenic cyanobacterial mats in rivers. We know of only one published study on this topic, which documented shifts over time in the microbial assemblages of *Phormidium* mats (Oscillatoriales) in New Zealand rivers (Brasell *et al.*, 2015). Proteobacteria were abundant in these assemblages, including taxa known to produce alkaline phosphatase, which may be important for mat growth in these phosphorus-limited New Zealand rivers (Wood *et al.*, 2015; McAllister *et al.*, 2016; Wood *et al.*, 2017). The New Zealand mats were profiled using 16S rRNA gene amplicons surveys (Brasell et al., 2015), a method that cannot provide specific information about metabolic functions or metabolites.

Cyanobacterial mats occur each summer in the Eel River in Northern California (Figure 1). The mats are formed by filamentous cyanobacterial taxa such as *Phormidium* (Oscillatoriales) and *Anabaena* (Nostocales) and produce cyanotoxins, especially anatoxin-a (Puschner *et al.*, 2008; Bouma-Gregson *et al.*, 2017). Anatoxin-a is a neurotoxic alkaloid that inhibits neuromuscular receptors by disrupting cellular ion channels, which causes muscle failure and can lead to death (Devlin and Edwards, 1977; Carmichael *et al.*, 1979). Anatoxin-a synthesis starts from proline, a cyclic amino acid, which is transformed by an eight-gene (*ana*A-H) polyketide synthase (PKS) operon (Méjean *et al.*, 2010, 2014; Cadel-Six *et al.*, 2009). Not all strains within anatoxin-a producing species contain the anatoxin-a gene operon, and variation between toxin and non-toxin producing strains occurs over small spatial scales (<1 cm) (Wood *et al.*, 2012; Wood and Puddick, 2017). Concentrations of anatoxin-a can be driven by changes in the abundance of toxin-producing genotypes within a mat, rather than differential toxin production per cell (Wood and Puddick, 2017).

**Figure 1.**
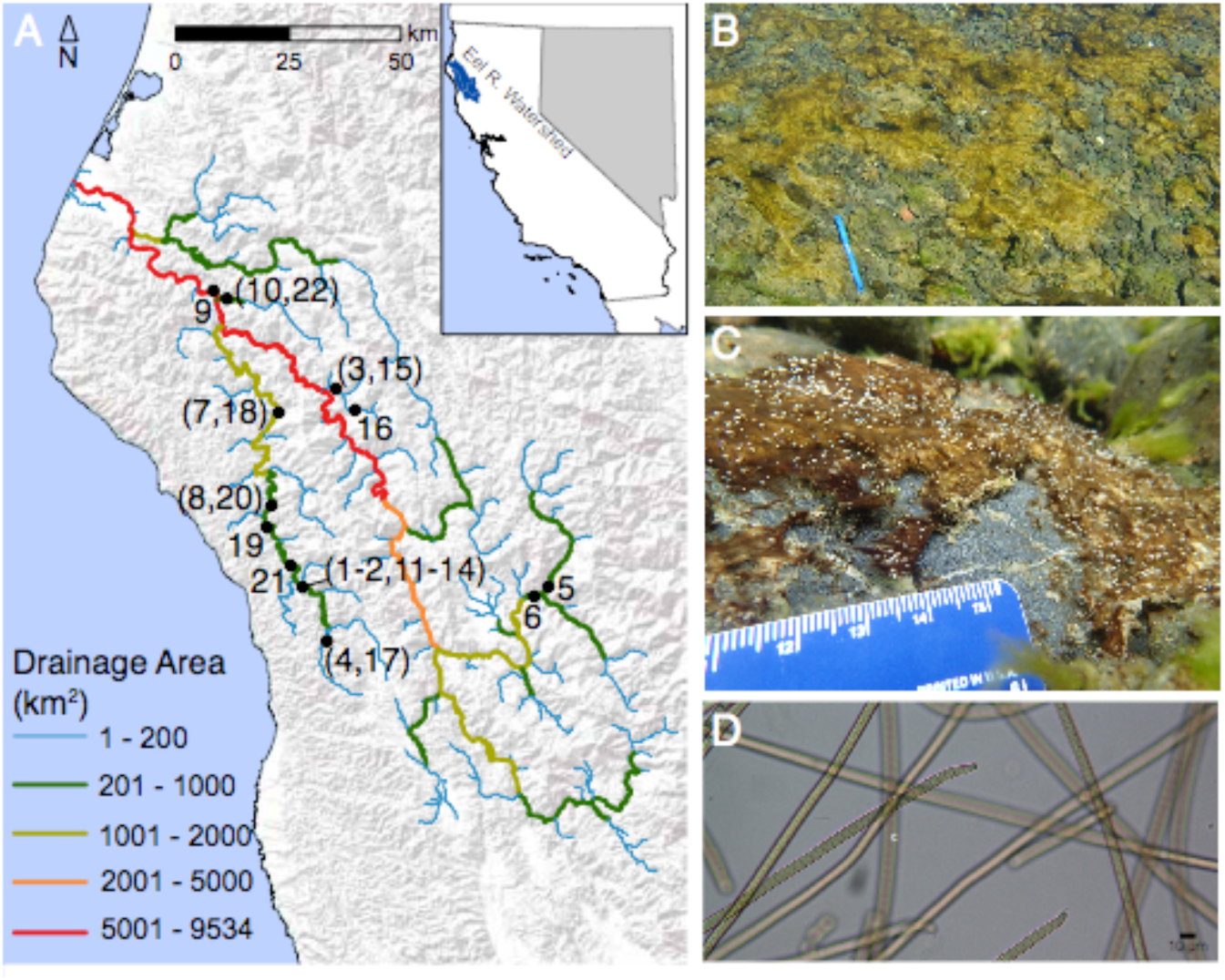
A) Map of Eel River watershed showing location of 22 samples collected in the Eel River watershed in August 2015. B) Phormidium mats in the Eel River. C) Underwater photograph showing *Phormidium* mat on a cobble. D) Micrograph of *Phormidium* trichomes (400x).

In spite of their important ecological and public health impacts, few toxin-producing cyanobacterial genomes have been sequenced (Shih *et al.*, 2013; Brown *et al.*, 2016; Pancrace *et al.*, 2017). Currently, there have been no culture-independent genome-resolved investigations of toxigenic freshwater cyanobacterial mats, which can provide high-resolution information about taxonomic diversity and provide direct insight into the metabolic potential of microbes. Using genome-resolved metagenomics, we studied *Phormidium*-dominated mats across the Eel River watershed to 1) identify the diversity of toxin-producing *Phormidium*, 2) document the diversity of non-cyanobacterial taxa associated with the *Phormidium* mats, 3) describe the anatoxin-a gene operon of *Phormidium* within the watershed, and 4) identify different metabolic potential of Bacteria present in mats.

## Materials and methods

### Sample collection

Samples were collected from *Phormidium* (Oscillatoriales) mats over 3 weeks in August 2015 in the Eel River watershed in California (Figure 1). The upstream drainage areas at sites ranged from 17 km^2^ to 7908 km^2^. *Phormidium* mats were identified macroscopically by their brown, orange-olive, or maroon coloration and epilithic growth. *Phormidium* mats are usually found in riffles, and all samples were collected from water 5-70 cm deep with surface flows of 5-100 cm s^-1^. Microscopic identification confirmed that the mats are dominated by *Phormidium* (Figure 1). At each site, a cobble covered by a *Phormidium* mat was lifted from the river and placed in a tray that had been cleaned using a 70% ethanol solution. Using sterile forceps, ~0.5 g of biomass was sampled from the mat and placed in a sterile 2 ml cryotube. Samples were immediately flash-frozen by filling a Styrofoam cooler with 2 L of ethanol and then adding ~250 ml of dry ice to rapidly lower the temperature. Cryotubes were immediately placed in a plastic bag and submerged in the ethanol for 2 minutes. Then cryotubes were stored on dry ice until placed at -80°C upon returning to the laboratory. When enough *Phormidium* biomass could be located at a site, an upstream and downstream sample was collected about 30-50 meters apart from each other. In total, 22 samples were analyzed from 13 sites (Figure 1).

Several environmental parameters were measured at each site: total dissolved nitrogen and phosphorus, nitrate, ammonium, depth, surface flow velocity, canopy cover, conductivity, temperature, dissolved oxygen, alkalinity, and pH (collection and analysis details in supplementary methods). The relationship of samples based on environmental variables was analyzed with principal components analysis using the vegan package (Oksanen *et al.*, 2017) in R v. 3.4.2 (R Core Team, 2017). After a mat sample was collected for DNA extraction, ~1 g of cyanobacterial mat remaining was collected to measure anatoxin-a using liquid-chromatography mass spectrometry (LC-MS) (details in supplementary methods).

### DNA extraction and sequencing

DNA was extracted from samples using a MoBio (Carlsbad, CA, USA) DNeasy PowerBiofilm kit. Frozen cyanobacterial mat samples were thawed at room temperature for 0.5 h, and approximately 0.15 g of mat removed for DNA extraction. The DNA extraction followed manufacturer’s protocol, except the cell lysis step in the protocol was modified to 5 minutes of bead beating and submersion for 30 minutes in a 65°C water bath. DNA was eluted into double distilled H_2_O, and sequenced on an Illumina (San Diego, CA, USA) HiSeq 4000 with 150 bp paired-end reads at the QB3 Genomics Sequencing Laboratory (http://qb3.berkeley.edu/gsl/, Berkeley, CA, USA).

### Metagenome assemblies, binning, and analyses

Reads were filtered to remove Illumina adapters and contaminants with BBtools then trimmed with SICKLE (https://github.com/najoshi/sickle) using default parameters. Assembly and scaffolding was performed by IDBA_UD (Peng et al., 2012). For assembled scaffolds longer than 1 kbp, protein-coding genes were predicted with Prodigal in the meta-mode (Hyatt et al., 2010). Predicted genes were then annotated against KEGG (Kanehisa et al., 2014), UniRef100 (Suzek et al., 2007), and UniProt using USEARCH (Edgar, 2010). Genomes were binned manually using coverage, GC content, single copy genes, and taxonomic profile with ggKbase (ggkbase.berkeley.edu), as described in Raveh-Sadka et al. (2015). Raw sequence data and assembled *Phormidium* genomes have been deposited to NCBI under BioProject number PRJNA448579.

The taxonomic composition of the microbial assemblage in samples was investigated using the ribosomal protein S3 (rpS3) gene, and average nucleotide identity (ANI) used to investigate diversity of *Phormidium* genomes (details in supplementary methods). In genomes >70% complete and with <10% contamination, according to CheckM (Parks *et al.*, 2015), genes for different metabolisms, phosphorus cycling, and anatoxin-a biosynthesis were predicted using the annotation process described above, as well as hidden Markov-models and read mapping (details in supplementary methods).

## Results

### Environmental conditions

At the collection sites, temperatures ranged from 17-26°C and maximum dissolved nitrogen and phosphorus concentrations were <100 and <30 µg L^−1^, respectively (Table S3). From the principal component analysis (PCA) on the environmental variables, the first two axes explain 60.7% of the environmental variation among the sites. Canopy cover and temperature were negatively correlated and explain much of the variation in environmental conditions at each site along PCA axis 1 (Figure S1). The second PCA axis describes variation in nitrogen, pH, and dissolved oxygen.

### Microbial assemblage diversity

The most abundant organisms in microbial assemblages belonged to Cyanobacteria (18 rpS3 clusters), Bacteroidetes (97 rpS3 clusters), Proteobacteria (77 rpS3 clusters), and Verrucomicrobia (8 rpS3 clusters). Minor members of the community included bacteria from Actinobacteria, Deinococcus-Thermus, Chloroflexi, *Canditatus* Kapabacteria and three Candidate Phyla Radiation groups (CPR; Figure S2). The genomic dataset for each sample included a relatively unique set of rpS3 sequence clusters, resulting in low species overlap (high beta diversity) among the sites with mean and median *β*sim values of 0.73 and 0.8. Additionally, 25% of pairwise comparisons among sites had *β*sim values of 1, indicating no shared rpS3 clusters among the samples. No single rpS3 cluster was detected in all samples.

Based on the relative abundance of rpS3 clusters, Cyanobacteria were the most abundant organisms in all samples, comprising 62-98% of the relative abundance within each sample (Figure 2A and Figure S3). The most common cyanobacterial rpS3 clusters were rpS3 66, 85, 152, and 244. These formed a clade with *Oscillatoria nigro-viridis* and *Microcoleus vaginatus* FGP-2 (Figure S2), but not with any reference *Phormidium* rpS3 sequences (Figure S2). (Few *Phormidium* rpS3 reference sequences are available, and the designations of the genera *Oscillatoria*, *Microcoleus*, and *Phormidium* are under revision (Struneck*ý et al.*, 2013; Komárek, 2017). Therefore, we refer to the dominant Oscillatoriales genomes in each sample as *Phormidium*). Bacteroidetes were the most abundant non-cyanobacterial taxa, occurring in all 22 samples, representing 35-100% of the non-cyanobacterial assemblage (Figure 2B). Proteobacteria, primarily Burkholderiales and Sphingobacteriales, were also common in samples. Verrucomicrobia were detected in 10 samples at relative abundances approaching >40% (Figure 2B). The four CPR rpS3 clusters that occurred at low abundance were identified as Peregrinibacteria (PER), Absconditabacteria (SR1) and Parcubacteria (OD1) (Figure S2).

**Figure 2.**
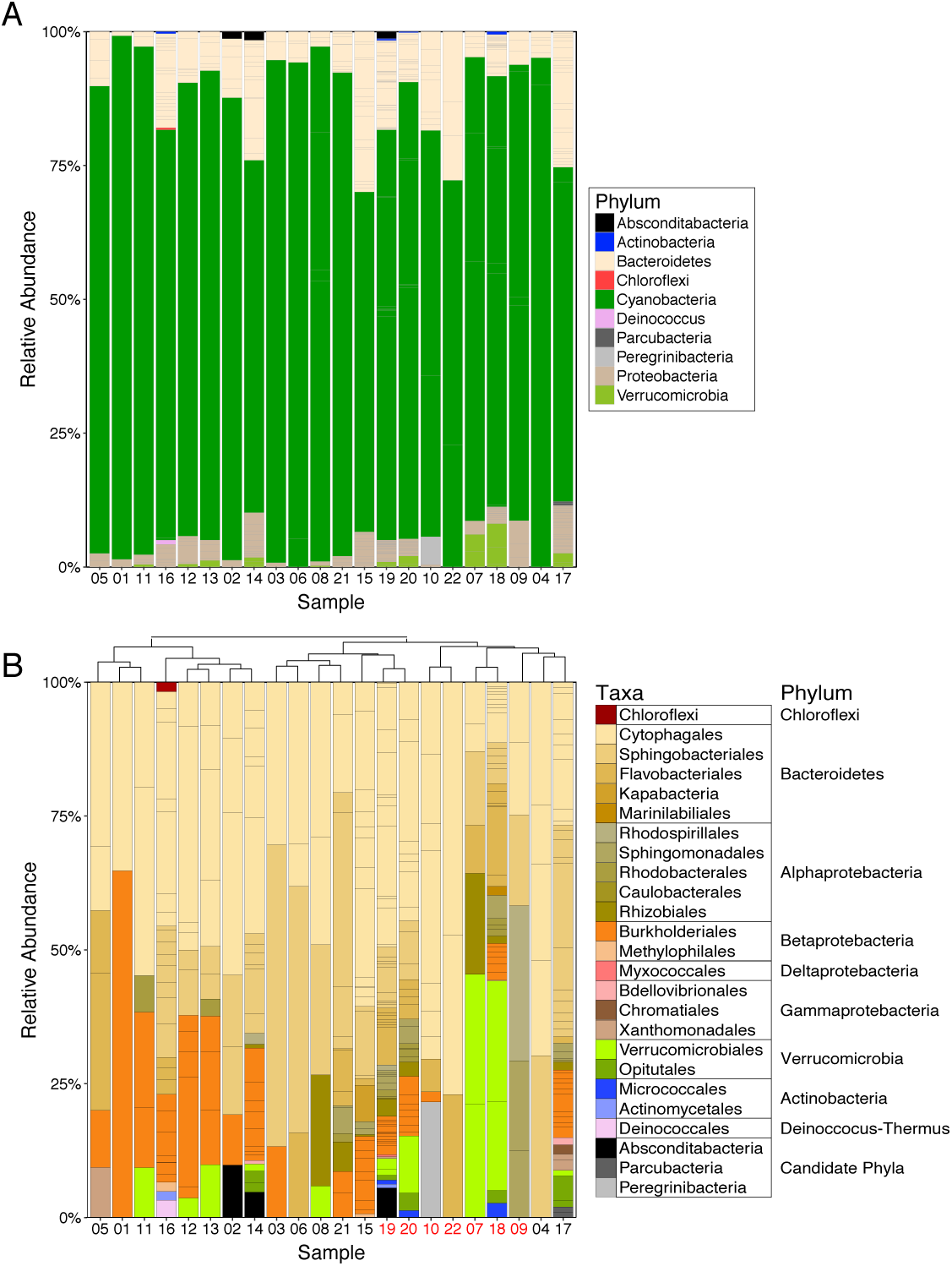
Percent relative abundance of rpS3 sequence clusters in samples. A) rpS3 clusters are colored by phyla. B) Non-cyanobacterial rpS3 clusters colored at a variety of different taxonomic levels that indicate the best classification given the bin novelty. Columns are clustered by Ward’s distance and samples in red indicate the recovery of the anatoxin-a gene operon in that sample.

Genomic analyses showed that each of the 22 mats was dominated by Cyanobacteria from the order Oscillatoriales, as represented by a single draft genome for a population of very closely related organisms. In 12 samples, a second or third *Phormidium* genome was also recovered at lower coverage. Interestingly, these genomes formed four *Phormidium* species clusters sharing less than 96% nucleotide identity (the species level ANI threshold) (Varghese *et al.*, 2015) among the clusters (Figure 3A). *Phormidium* species 1-3 had 82-88% ANI with reference genomes. However, *Phormidium* species 4 had 96.1% ANI with the *Phormidium willei* BDU130971 reference genome, suggesting these might be the same species. In samples with more than one recovered *Phormidium* genome, the two genomes belonged to different species, with the second genome occurring at much lower coverage (Figure 3A). Typically one or two rpS3 sequence clusters were associated with each *Phormidium* species.

**Figure 3.**
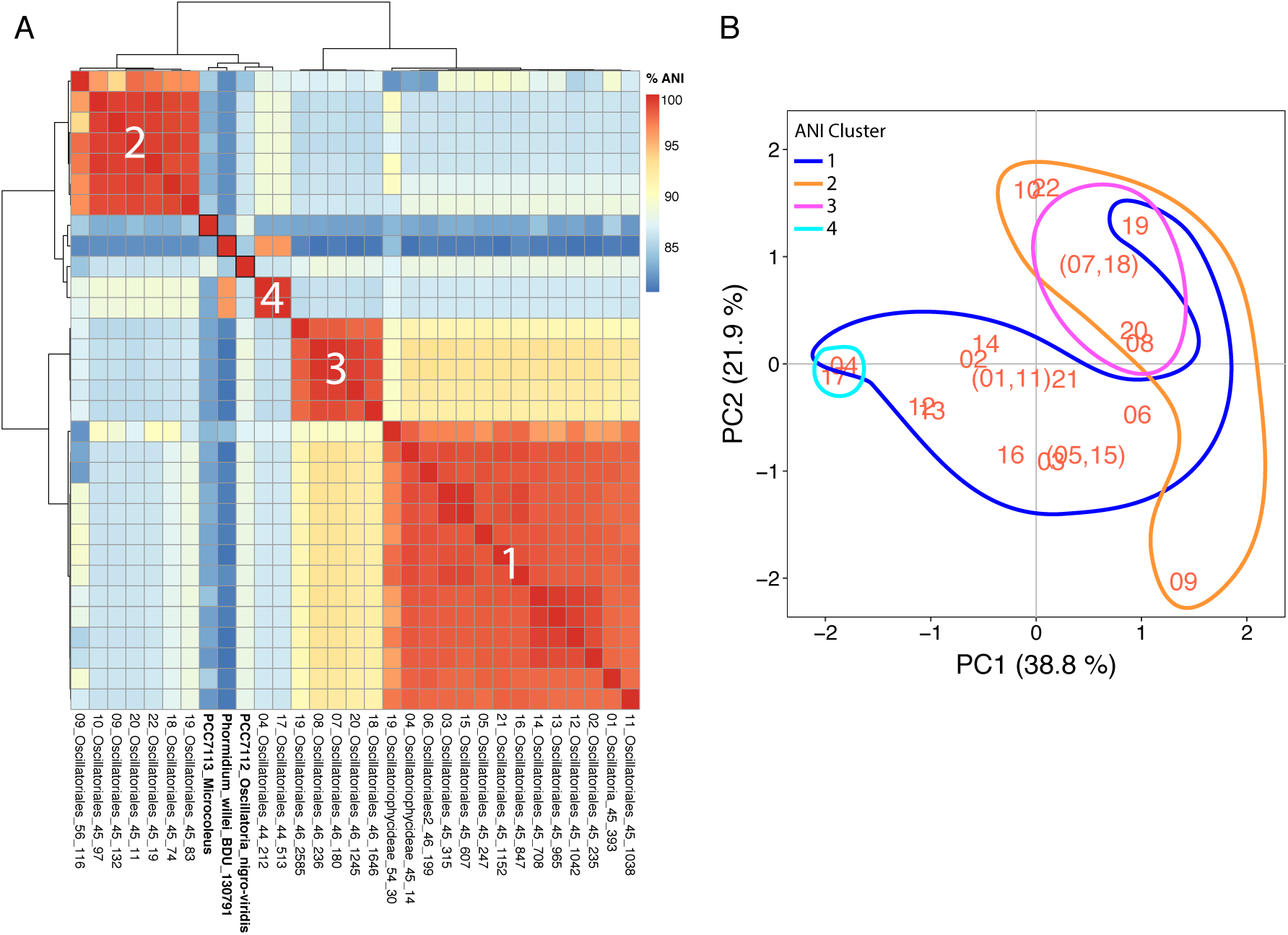
A) Percent average nucleotide identity (ANI) of cyanobacterial genomes from samples and three reference genomes (in bold). The first number of the genome name identifies the sample location (See Figure 1). The latter two numbers indicate the GC content and genome coverage. ANI results were clustered using Ward’s distance and represent 4 species groups with <93% ANI among the clusters. B) Principal components axes of environmental variables from Figure S1 with colored lines surrounding samples from each ANI cluster. Some samples generated genomes from multiple ANI clusters.

With the exception of *Phormidium* species 1, *Phormidium* species were limited to sub-regions in the watershed. Species 1 contained genomes collected from sites broadly distributed across the watershed (Figures 1 and 3B), including the eastern sites 05-06 and 15-16. However, species 1 also contained a sub-cluster of samples 02 and 12-14 with 99-100% ANI, all collected within 0.5 km in a single creek. Species 2 was found in the most downstream region (samples 09, 10, and 22) and in the middle reaches of the South Fork Eel (samples 18-20). Species 3 was limited to 3 sites in the South Fork Eel (samples 07-08 and 18-20), and Species 4 came from a single site (samples 04 and 17). Each *Phormidium* species also came from a wide range of environmental conditions, with only Species 3 and 4 found in a smaller environmental niche (Figure 3B).

### Anatoxin-a

Metagenomic assembly allowed the recovery of the anatoxin-a synthesis gene operon (*ana*A a– *ana*J) from 7 samples: 07, 09, 10, 18, 19, 20, and 22 (Figure 4A). Read mapping to the anatoxin operon in sample 19 (scaffold PH2015_19_scaffold_1561) did not detect any anatoxin synthesis genes in other samples (Table S4). The non-cyanobacterial microbial assemblage was different in samples with and without the ATX operon (Figure 4C, p <0.01). This relationship is primarily driven by fewer Betaproteobacteria in samples without the ATX operon (Figure 4B, p <0.01).

**Figure 4.**
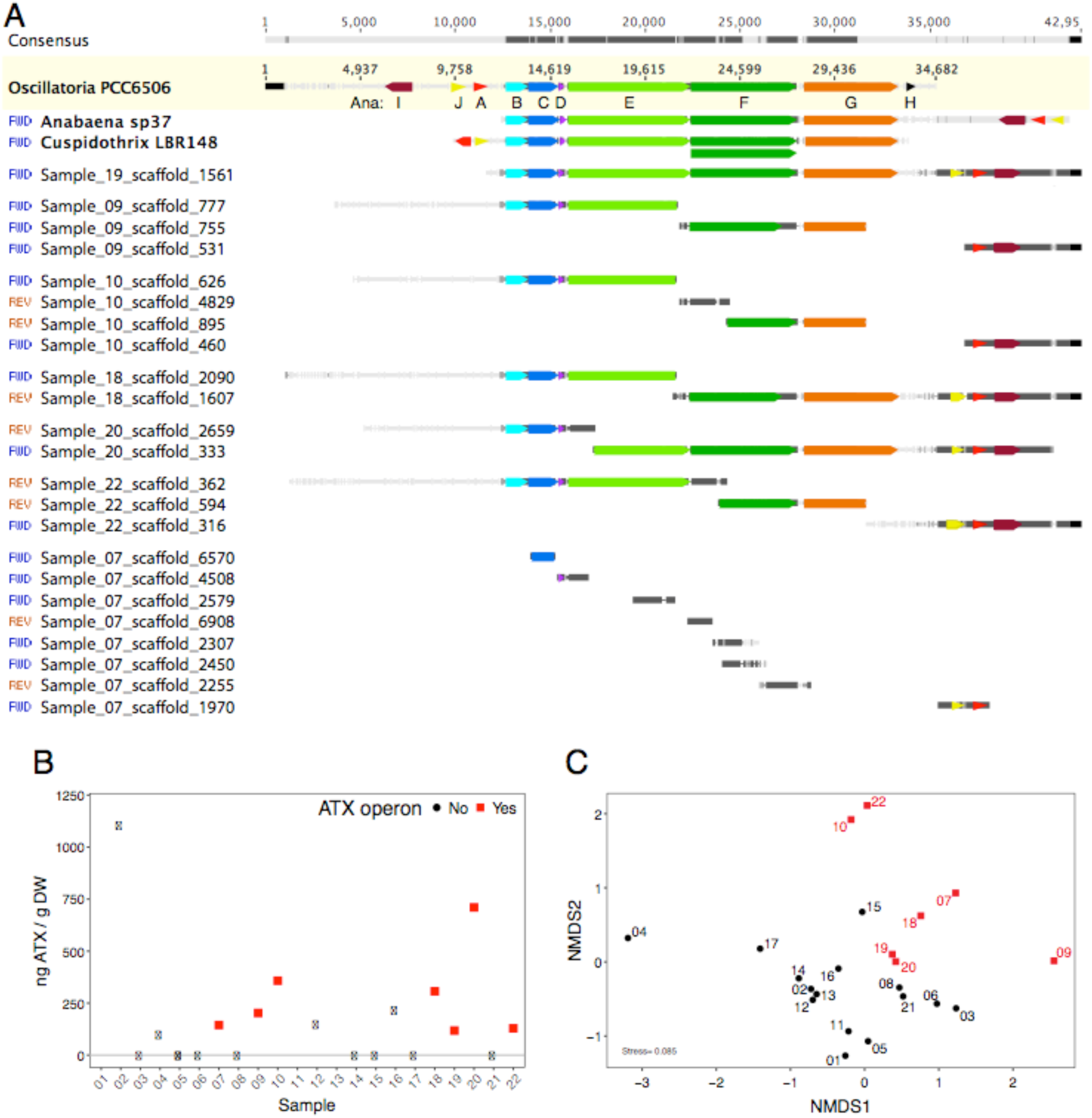
A) Anatoxin-a operon from three reference sequences (bold) and samples 19, 09 10, 18, 20, 22, and 07. Sample scaffolds were mapped to Oscillatoria PCC6505, and gene annotations added at 50% identity to one of the 3 reference sequences. Different *ana* genes are different colors. B) Anatoxin-a (ATX) concentrations in *Phormidium* mat samples. ATX was not measured in samples 01, 11, and 13. C) Non-metric multidimensional scaling (NMDS) plot using Bray-Curtis dissimilarities of non-cyanobacterial assemblage showing samples with (red squares) and without (black circles) the anatoxin-a operon.

LC-MS analyses detected anatoxin-a in 11 of the 19 samples (Figure 4B), with median and maximum concentrations of 153 and 1104 ng anatoxin-a / g DW, respectively. In four samples, anatoxin-a was detected with the LC-MS, but the operon was not recovered. There was no difference in the microbial assemblage among samples with and without LC-MS detection of anatoxin-a (p = 0.089).

In all samples but 07, the operons were found in the rpS3_152 bin and in *Phormidium* species 2 (Figures 3 and 5A). In sample 07, the operon was partially recovered from the rpS3_85 bin in *Phormidium* species 3, with the *ana*C, J, and I genes assembled and annotated. Other scaffolds in sample 07 mapped to the PCC6506 operon, but the genes did not annotate to the assembled scaffolds at >50% identity. Most nucleotide identities of the *ana* genes to the PCC6506 reference sequence ranged from 88-94% similar with lower values in the *ana*F, G, and J genes (Table S5). Amino acid *ana*C sequences were conserved among the samples (100% identity), but other genes in the operons were more variable (*ana*E, F, G, and J; Table S5).

**Figure 5.**
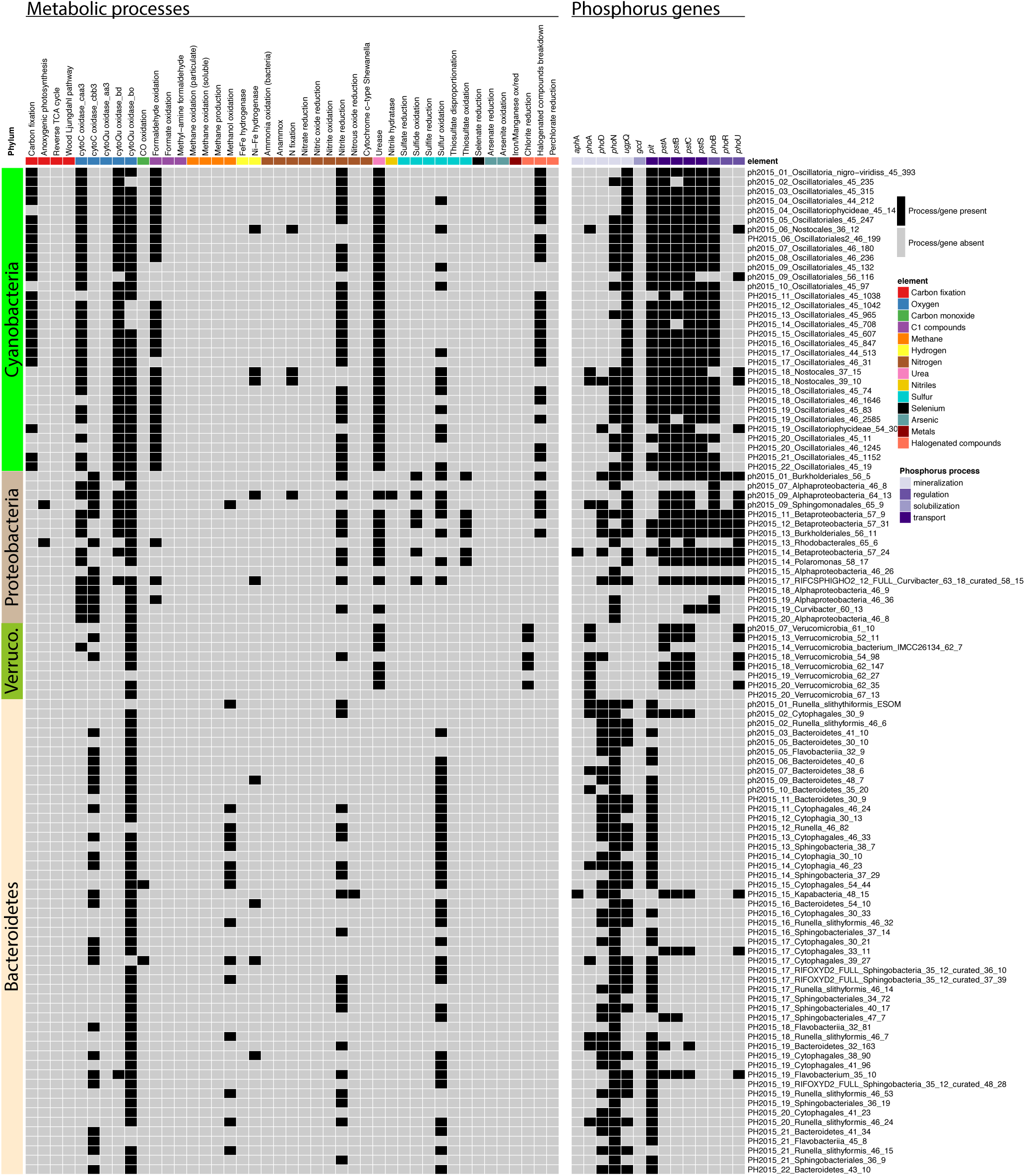
Heatmap showing the presence or absence of different metabolic processes (left panel) and phosphorus acquisition and transport genes (right panel) in reconstructed genomes. Each genome is a row and the genome name is listed on the right. Each metabolic or phosphorus processes is a column. Column colors indicate different metabolism types (left panel), or genes involved in phosphorus acquisition and transport. A process or gene that is predicted in a genome is indicated by a black square. The phyla of the genomes is listed to the left of the heatmap.

All anatoxin-a gene annotations from these sequences matched those previously reported (Rantala-Ylinen *et al.*, 2011; Méjean *et al.*, 2014; Jiang *et al.*, 2015; Brown *et al.*, 2016).

The arrangement of the *ana*B-G genes was similar to the three reference operons (Figure 4A). However *ana*A, I, and J genes were located on the opposite end of the operon compared to PCC6506, and closer to *ana*G than the two Nostocales reference operons (*Anabaena* and *Cuspidothrix*) (Figure 4A). In samples 09 and 10, *ana*J was not assembled, but read mapping confirmed its presence in these samples. The transposase *ana*H gene was not found in any samples. The *ana*G gene was ~1670 base pairs shorter than in the three reference sequences. This missing region contained the ~300 base pair methyltransferase domain, proposed to affect the production of either anatoxin-a of homoanatoxin-a (Méjean *et al.*, 2014, 2016). All samples also lacked the recently identified *ana*K gene, which enables production of dihydroanatoxin-a (Méjean *et al.*, 2016).

### Metabolic potential and phosphorus acquisition

Considering the entire microbial community, the most abundant metabolisms across samples were phototrophic, heterotrophic, nitrogen, and sulfur metabolisms (Figure 5). Although genes associated with nitrogen fixation, hydrogen oxidation, halogenated compounds, and formaldehyde were also detected, the most common processes were carbon fixation, oxygen respiration, nitrite reduction, utilization of urea, and sulfur oxidation. In addition to carbon fixation, *Phormidium* genomes contained genes for formaldehyde oxidation, urea breakdown, nitrite reduction, sulfur oxidation, and halogenated compounds breakdown. Across the 4 *Phormidium* species, there was little variation in metabolic potential.

Proteobacteria had the most diverse sulfur metabolic potential in the mats. Genes for sulfide oxidation (flavocytochrome c sulfide dehydrogenase), sulfur oxidation (sulfur dioxygenase), and thiosulfate oxidation (*sox*B, C, Y) were identified in genomes from both Alpha and Betaproteobacteria. Anoxygenic photosynthesis genes were found in a Sphingomonadales and a Rhodobacterales genome. A phylogeny of rpS3 sequences shows that the PH2015_09_Sphingomonadales_65_9 genome clusters with the freshwater aerobic anoxygenic phototrophic bacteria *Porphyrobacter* spp. and the PH2015_13_Rhodobacterales_65_6 genome with the purple non-sulfur bacteria Rhodobacterales sphaeroides and Rhodobacterales capsulatus (Figure S4). Aerobic anoxygenic phototrophic bacteria are often found in eutrophic sunlit aquatic environments and are heterotrophs that use anoxygenic photosynthesis to provide supplemental energy for cell growth and maintenance (Yurkov and Beatty, 1998; Koblizek, 2015).

Organisms in the phyla Bacteroidetes had genes for few metabolic processes. Based on analyses of 50 Bacteroidetes genomes, most genomes contain genes for carbon and sulfur oxidation (Figure 5). However, Bacteroidetes were the only organisms predicted to possess genes for oxidation of methanol to formaldehyde. Verrucomicrobia primarily have genes for carbon oxidation and nitrite reduction. Bacteria from this phylum are the only organisms found in the mats with predicted genes for chlorite reduction (Figure 5).

Phosphorus concentrations were low at most sites (<10 µg L^−1^; Table S3), and phosphorus uptake and transport genes were present in most genomes (Figure 5). The non-cyanobacterial assemblage contains genes for alkaline phosphatase, while the most common phosphorus mineralization genes for *Phormidium* were acid phosphatase *pho*N and glycerophosphodiesterase *ugp*Q. The low-affinity inorganic phosphorus transport gene (*pit*) was most frequently identified in the Cyanobacteria and Bacteroidetes genomes, but all Verrucomicrobia and most Proteobacteria lacked this gene. *Phormidium* also contained the high affinity phosphate transport pst genes. Unlike other phyla, Verrucomicrobia only possessed *pho*A and the *pst* transporter genes. The phosphorus-solubilization gene, gcd, was absent in all genomes.

## Discussion

This is the first genome-resolved metagenomic analysis of toxin-producing freshwater benthic cyanobacterial mats. By recovering the genomes of bacteria within 22 *Phormidium* mats collected across a 9547 km^2^ watershed, we provide information about the microbes and genes potentially controlling energy flows, nutrient cycling, and toxin production within these benthic *Phormidium* mats.

### Microbial diversity

The dominant phyla in our samples, Cyanobacteria, Bacteroidetes, Proteobacteria, and Verrucomicrobia, have been found to co-occur with freshwater planktonic cyanobacterial blooms (Pope and Patel, 2008; Steffen *et al.*, 2012; Mou *et al.*, 2013; Louati *et al.*, 2015; Parulekar *et al.*, 2017), and were also abundant in riverine *Phormidium* mat assemblages in New Zealand (Brasell *et al.*, 2015). Prior studies of planktonic cyanobacterial blooms found a higher relative abundance of Proteobacteria, compared to Bacteroidetes, than we did in our benthic samples. In most prior studies, Alphaproteobacteria were the dominant group of Proteobacteria; only Louati et al. (2015) reported a higher relative abundance of Betaproteobacteria, as in our results. Increases in cyanobacterial biomass can affect microbial assemblages. As planktonic cyanobacteria blooms, or benthic *Phormidium* mats develop and thicken, the co-occurring microbial assemblage composition changes (Li *et al.*, 2012; Parveen *et al.*, 2013; Brasell *et al.*, 2015). This successional change could account for some of the variation observed within and across sites and ecosystems.

In contrast to the non-cyanobacterial species, the *Phormidium* species showed higher overlap among the sites, and environmental conditions were not strongly associated with the presence of specific groups within each sample (Figure 5). *Phormidium* species 4 was the exception, being found only in the coolest shadiest site. More sampling of smaller tributaries could test the hypothesis that *Phormidium* species 4 is a cool-water, low sunlight *Phormidium* species, while *Phormidium* species 1-3 tend to occur in warmer sunnier environments. All four assembled *Phormidium* species in this study had low sequence similarity (ANI) to reference genomes, suggesting each is a novel species. Each draft *Phormidium* genome probably represents a new species of Cyanobacteria, thus helps to fill important gaps in our understanding of cyanobacterial diversity and physiology.

### Microbial assemblage and anatoxin-a

The difference in the non-cyanobacterial assemblages between samples with and without the anatoxin-a operon suggests a relationship among anatoxin-a producing cyanobacteria and the non-cyanobacterial assemblage. Although it is possible that non-cyanobacterial organisms present in the environment influence selection for cyanobacterial strains, we consider the reverse more likely. As cyanobacteria bloom they will change environmental conditions, in part due to cyanobacterially-produced compounds, which then shape microbial community composition. This could involve selection for bacteria with the capacity to degrade cyanotoxins. Microcystin concentrations have been shown to affect bacterial assemblage composition (Mou *et al.*, 2013), and the copies of microcystin degradation genes (mlr) in the overall community have been positively correlated with microcystin concentrations (Li *et al.*, 2015; Lezcano, 2016).

Little is known, however, about anatoxin-a degrading bacteria (Kormas and Lymperopoulou, 2013). Anatoxin-a degradation rates are increased by Bacteria in sediments (Rapala *et al.*, 1994), and a *Pseudomonas* sp. capable of degrading anatoxin-a has been isolated (Kiviranta *et al.*, 1991). Mou et al. (2013) found a positive association between microcystin degradation rates and Burkholderiales, suggesting a role for these bacteria in toxin breakdown. Burkholderiales are important in some cyanobacterial blooms (Louati *et al.*, 2015), although a decline in abundance of a Burkholderiales strain can be predictive of the onset of a cyanobacterial bloom (Tromas *et al.*, 2017). In the current study, samples with the anatoxin-a operon had fewer Burkholderiales, so it possible that the bacteria from this family in the Eel River mats are negatively impacted by toxin production (unfortunately, the genes for anatoxin-a degradation are not known, so this cannot be determined genomically).

With the exception of samples 9 and 10, the other sample assemblages containing Cyanobacteria with the anatoxin-a operon had relatively more Sphingomonadales, an order with many known microcystin degrading species (Kormas and Lymperopoulou, 2013; Mou et al., 2013; Briand et al., 2016). However, anatoxin-a and microcystin have different molecular structures. Anatoxin-a is bicyclic alkaloid, while microcystin is a cyclic peptide. It cannot be assumed that genes involved in microcystin degradation will confer the ability to degrade anatoxin-a. Our findings motivate experiments to test impacts of co-occurring microbes on anatoxin-a degradation.

### Microbial assemblage and metabolisms

Strong redox gradients occur in microbial mats in salt marshes, lakes and mudflats and result in diverse spatially structured metabolisms (Armitage *et al.*, 2012; Franks and Stolz, 2009). Although the epilithic *Phormidium* mats from this study clearly derive their energy from photosynthesis and carbon oxidation in an aerobic environment, a small fraction of bacteria in the assemblages appear to derive energy using other metabolic pathways. However, the epilithic growth on larger cobbles, compared to sand or silt substrates, limits the depth of the mat and likely prevents development of the permanent anoxic regions found in cyanobacterial dominated mats in estuaries (Franks and Stolz, 2009). Within the Eel River mats, oxygen concentrations likely fluctuate on a diel cycle as they do in New Zealand (Wood *et al.*, 2015), creating transient anoxic conditions (Paerl and Pinckney, 1996) and limiting the relative abundance of organisms with anaerobic metabolisms. Overall, the mats studied here are metabolically simple, with few anaerobic or non-carbon metabolisms. This simplifies predictions about the environments in the river network in which mats may proliferate.

Genes for carbon respiration were common among reconstructed genomes from the mats, especially within Bacteroidetes genomes, the most abundant non-cyanobacteria in most samples. Bacteria from this phylum are sometimes heterotrophs, capable of breaking down complex organic compounds (Grondin *et al.*, 2017). Consistent with this prediction, many of the genomes encode large inventories of genes for degradation of complex carbohydrates (e.g., diverse glycosyl hydrolases). *Phormidium* mats are characterized by thick extracellular polymeric substances (EPS) that give structural integrity to the mat (Nicolaus *et al.*, 1999). Therefore, it is likely that Bacteroidetes are metabolizing carbon compounds associated with the EPS produced by *Phormidium*, (Beraldi-Campesi *et al.*, 2012).

Bacteroidetes were the only bacteria predicted to possess methanol oxidation genes, which oxidize methanol to formaldehyde. It is possible that some Bacteroidetes supply formaldehyde for *Phormidium* to metabolize to formate. This further suggests a close carbon association of Bacteroidetes with *Phormidium*, as most *Phormidium* genomes contain formaldehyde oxidation genes. However, as no formate metabolism genes were found, so the fate of formate is unclear.

Consistent with prior studies (Niemi *et al.*, 2009), Burkholderiales are predicted to have some capacity to degrade complex organic compounds. This may also explain their high abundance in mats. CPR bacteria may also contribute to carbon cycling. As noted previously (e.g., Kantor et al., 2013; Nelson and Stegen, 2015), the genomes of the bacteria from the three CPR groups are small and the organisms are predicted to have symbiotic anaerobic, fermentation-based lifestyles. Parcubacteria have been reported in planktonic cyanobacterial blooms (Parulekar et al., 2017), Doudnabacteria (SM2F11), and 12 other Candidate Phyla reported in Antarctic Cyanobacteria/diatom mats (Stanish et al., 2013), and Absconditabacteria (SR1) reported from various anaerobic aquatic habitats (Davis et al., 2009). Their identification in mats in a river network suggests that they play important and previously unrecognized roles in aquatic biofilms, both via carbon turnover and impacts on host organisms.

Many Cyanobacteria and Proteobacteria contain genes for sulfur and sulfide oxidation, both of which can occur in oxygenated environments, but often in proximity to anoxic regions where reduced sulfur compounds are formed or accumulate (Wasmund et al., 2017). However, most rivers do not have high concentrations of elemental or reduced sulfur in the water column (Meybeck, 1993). Therefore, it is likely that these organisms are not using sulfur as a primary energy source, but are capable of metabolizing sulfur opportunistically when it may sporadically accumulate in anoxic microhabitats in the mats.

In the Eel River, nitrogen is considered a limiting nutrient in spring and early summer (Hill and Knight, 1988; Finlay *et al.*, 2011). For example, epilithic periphyton assemblages are often dominated by nitrogen fixing taxa (Power *et al.*, 2009), such as endosymbiotic nitrogen fixing cyanobacteria associated with Rhopalodiaceae diatoms, *Nostoc* spp., or *Anabaena* spp. However, few nitrogen fixation genes were predicted in the dataset analyzed here. Though some *Phormidium* species can fix nitrogen (Bergman *et al.*, 1997), as expected based on samples in New Zealand (Heath *et al.*, 2016), the *Phormidium* in Eel River mats do not fix nitrogen. Nitrogen fixing capacity was found in a single Alphaproteobacteria genome and three Nostocales genomes. *Phormidium* and most other bacteria in the Eel River mats likely derive their nitrogen exogenously from a combination of the water column and Rhopalodiaceae diatoms and the few nitrogen fixing bacteria living in the mats.

All *Phormidium* genomes, and most Proteobacteria and Verrucomicrobia genomes, are predicted to contain urease genes. Dissolved organic nitrogen is hypothesized to be an important nitrogen source in the summer in the Eel River (Finlay *et al.*, 2011). Urease genes may also suggest high rates of nitrogen cycling within mats, as organic nitrogenous waste products from organisms are taken up by other organisms. Therefore, once mats establish, internal cycling of organic nitrogen may decouple the mats from nitrogen dynamics in the overlying water column (Vadeboncoeur and Power, 2017).

Phormidium mats proliferate in New Zealand rivers at low dissolved phosphorus concentrations (McAllister *et al.*, 2016; Wood *et al.*, 2017), and phosphorus may become increasing limiting in the Eel River in late summer (Finlay *et al.*, 2011), when *Phormidium* mats are more commonly observed. Based on our genomic analyses, many of the organisms in the microbial assemblage likely contribute to phosphorus mineralization in mats. The pH levels within *Phormidium* mats in New Zealand oscillate daily due to changes in CO_2_ and O_2_ concentrations driven by cyanobacterial photosynthesis (Wood *et al.*, 2015). *Phormidium* may use acid phosphatases to scavenge phosphorus at night when pH levels drop, and non-cyanobacteria use alkaline phosphatases scavenge during the day when pH is elevated. In New Zealand, phosphate weakly sorbed onto sediments trapped in mats may be an important phosphorus source to *Phormidium* mats (Wood *et al.*, 2015). The lack of the solubilization gcd gene suggests that microbes are relying on pH and O_2_ gradients rather than extracellular enzymes to solubilize sorbed phosphorus. *Phormidium*, and some Proteobacteria, also contain the transporter pst genes, which are up-regulated at low inorganic phosphorus concentrations (Hirota *et al.*, 2010). The presence of phosphatase genes and pst transporter genes may enable *Phormidium* strains to outcompete other organisms for phosphorus and enable them to dominate periphyton assemblages at low phosphorus concentrations.

## Conclusion

*De novo* genome assembly of metagenomic datasets provided information about the community membership of *Phormidium*-dominated cyanobacterial mats in a river network and the metabolic capacities of the abundant Bacteria. Thus, it was possible to describe likely ecological exchanges of nutrients and energy in these mats. Similar to planktonic cyanobacterial blooms, but in contrast to laminated cyanobacterial mats on finer substrates, riverine mat metabolisms are primarily fueled by phototrophy and carbon degradation. Importantly, we directly identified the genes for cyanotoxin production and community compositional features that may predict the presence of this capacity. As humans put more pressure on water resources, through water extraction, nutrient pollution, and warming temperatures, cyanobacterial blooms are predicted to have increasing impacts on freshwater resources (Paerl *et al.*, 2011; Taranu *et al.*, 2015; Monteagudo and Moreno, 2016). Understanding the relationships among toxigenic cyanobacteria and abiotic and biotic factors will be necessary to predict or mitigate cyanobacterial blooms in the future.

## Acknowledgements

Housing and laboratory facilities were provided by the UCNRS Angelo Coast Range Reserve. We would like to thank the Reserve Manager, Peter Steel, for his assistance. Addien Wray, Natalie Soto, Lindsey Bouma-Gregson, and Brian and Wendy Gregson assisted with sample collection in the field. Sue Spalding offered DNA extraction guidance, and Arianna Nuri assisted with DNA extractions and other lab work. We thank Raphael Kudela for measuring anatoxin-a concentrations. Support was provided by the National Science Foundation’s Eel River Critical Zone Observatory [EAR-1331940] and a Department of Energy grant [DOE-SC10010566]. KBG was supported by a US Environmental Protection Agency STAR Fellowship [91767101-0]. AJP was supported by the German Science Foundation [DFG PR 1603/1-1]. This work used the Vincent J. Coates Genomics Sequencing Laboratory at UC Berkeley, supported by an NIH Instrumentation Grant [S10 OD018174].

